# *BrainUnit*: Integrating Physical Units into High-Performance AI-Driven Scientific Computing

**DOI:** 10.1101/2024.09.20.614111

**Authors:** Chaoming Wang, Sichao He, Shouwei Luo, Yuxiang Huan, Si Wu

**Affiliations:** School of Psychological and Cognitive Sciences, Peking University, Beijing, China; PKU-IDG/McGovern Institute for Brain Research, Peking University, Beijing, China; Center of Quantitative Biology, Peking-Tsinghua Center for Life Sciences, Academy for Advanced Interdisciplinary Studies, Peking University, Beijing, China; School of Mathematical Sciences, Shanghai Jiao Tong University, Shanghai, China; Institute of Natural Sciences, Shanghai Jiao Tong University, Shanghai, China; Guangdong Institute of Intelligence Science and Technology, Guangdong, China

## Abstract

Artificial intelligence (AI) is revolutionizing scientific research across various disciplines. The foundation of scientific research lies in rigorous scientific computing based on standardized physical units. However, current mainstream high-performance numerical computing libraries for AI generally lack native support for physical units, significantly impeding the integration of AI methodologies into scientific research. To fill this gap, we introduce `BrainUnit`, a unit system designed to seamlessly integrate physical units into AI libraries, with a focus on compatibility with JAX. `BrainUnit` offers a comprehensive library of over 2000 physical units and more than 300 unit-aware mathematical functions. It is fully compatible with JAX transformations, allowing for automatic differentiation, just-in-time compilation, vectorization, and parallelization while maintaining unit consistency. We demonstrate `BrainUnit`’s efficacy through several use cases in brain dynamics modeling, including detailed biophysical neuron simulations, multiscale brain network modeling, neuronal activity fitting, and cognitive task training. Our results show that `BrainUnit` enhances the accuracy, reliability, and interpretability of scientific computations across scales, from ion channels to whole-brain networks, without impacting performance. By bridging the gap between abstract computational frameworks and physical units, `BrainUnit` represents a crucial step towards more robust and physically grounded AI-driven scientific computing.

## 1 Introduction

Artificial intelligence (AI) technologies are revolutionizing the landscape of scientific research in unprecedented ways, profoundly impacting fields such as physics [36], chemistry [8], biology [2], climate science [20], astronomy [28], and neuroscience [31]. AI assists scientists in formulating hypotheses, designing experiments, efficiently processing and analyzing vast datasets, and uncovering insights that would be difficult or impossible to achieve with traditional methods alone [41, 43]. At the heart of Artificial Intelligence for Science (AI4S) is the powerful computational infrastructure provided by high-performance computing (HPC) libraries. Libraries like PyTorch [30], TensorFlow [1], and JAX [10] offer the computational efficiency, flexibility, and hardware acceleration necessary to meet the rigorous demands of AI-driven research. However, despite their strengths, these libraries were originally developed for machine learning and deep learning applications, and generally lack critical features needed for scientific research, such as the proper handling of physical units.

A fundamental pillar of scientific research is the precise measurement of the physical world, which relies on well-defined unit systems [14]. For example, the metric system, with the meter as the standard unit for length, ensures the interoperability and comparability of measurement results across different studies. Without such unified standards, effective communication and comparison of scientific results would be nearly impossible. In the realm of scientific computing, the accurate application of physical units is not only a mark of scientific rigor but also essential for preventing dimensional errors and avoiding catastrophic failures, such as the Mars Climate Orbiter disaster caused by unit mismanagement [25]. Moreover, maintaining a consistent unit system throughout complex computations is vital for reinforcing physical constraints in models and ensuring that results remain physically plausible. Standardized unit systems, such as the International System of Units (SI) [4], are invaluable for fostering global scientific collaboration, facilitating data sharing, and ensuring the accurate interpretation and reproducibility of research findings.

Despite their critical importance, current mainstream HPC libraries generally lack native support for physical units. This deficiency presents significant challenges for researchers attempting to apply advanced AI technologies to scientific discovery. In complex computational processes, even minor errors in unit management can lead to significant deviations in results. Consequently, researchers often resort to manual unit conversions, which not only increases operational complexity and the risk of errors but also consumes considerable time and effort that could otherwise be devoted to core research activities. Moreover, code that omits unit information is often difficult to interpret, leading to “magic numbers” that lack clear physical meaning. When computational anomalies occur, the absence of unit information significantly complicates troubleshooting and debugging. Furthermore, varying unit conventions across different research fields increase communication costs and create barriers to data and model sharing, hindering the cross-pollination of ideas that is often essential for groundbreaking discoveries [3]. Therefore, as the scientific community increasingly embraces AI-driven methodologies [41, 43], the need for standardized and comprehensive unit support within HPC libraries has become more urgent than ever.

To address this critical gap, we introduce `BrainUnit`, a unit system designed to seamlessly integrate physical units into HPC numerical libraries, with a particular focus on JAX [10]. `BrainUnit` seeks to transform scientific computing by marrying the computational power of modern AI libraries with the rigorous physical unit constraints that are fundamental to scientific inquiry. It provides a robust, efficient, and user-friendly framework for managing physical units, reducing potential errors, enhancing research productivity, and establishing unit standards. We delve into the design principles and implementation details of `BrainUnit` and its potential applications on brain dynamics modeling, illustrating how it can bridge the gap between the abstract world of high-performance computing and the physical realities that scientists strive to understand. As we stand on the brink of a new era in AI-driven scientific discovery [41, 43], `BrainUnit` represents a pivotal step towards fully harnessing the potential of AI to advance human knowledge and understanding.

### 1.1 Related Works

In the programming ecosystem of scientific computing, we already have excellent physical unit libraries, such as `Pint` and `Quantities` in Python, `Unitful.jl` in Julia, `Boost.Units` in C++, and `uom` in Rust. These libraries enable seamless operations, conversions, and calculations among physical units, greatly simplifying the processing and analysis of physical quantities for researchers. However, they face significant limitations. Notably, they lack compatibility with mainstream HPC frameworks like PyTorch [30], TensorFlow [1], and JAX [10], hindering their ability to meet the demands of complex scientific computations in parallel computing hardware such as GPUs and TPUs. Moreover, they generally employ the float64 data type for numerical representation. In contrast, AI4S applications, particularly those using neural networks, often require lower-precision data types to reduce memory usage and improve computational efficiency. This discrepancy may lead to unnecessary performance overhead and limit potential applications in resource-constrained scenarios. Additionally, these unit libraries typically lack support for flexible scaling, a crucial feature when dealing with physical quantities exhibiting extreme scale differences. In certain scientific research domains, improper handling of such units may result in precision loss, potentially compromising the accuracy and reliability of research outcomes. What’s more, a critical limitation of current physical unit systems is their lack of support for automatic differentiation (AD), which is fundamental to modern AI models [9]. This deficiency makes high-order differentiable optimization particularly challenging, limiting the integration of physical units into modern AI-driven research and development pipelines. Therefore, although existing works have made significant strides in advancing the unit standardization in scientific computing, we still need to continuously explore solutions to achieve deep integration with HPC frameworks.

### 1.2 Design Principles

Towards a standardized physical unit system, `BrainUnit` was designed and implemented with several key considerations:

1. ***Standardization***: Adopting SI Units to ensure global scientific compatibility and standardized communication.
2. ***Consistency***: Maintaining strict adherence to physical laws in the relationship between fundamental units (e.g., length, time, mass) and compound units (e.g., velocity, force, energy), ensuring logical consistency throughout the system.
3. ***Universality***: Comprehensive support for known physical quantities, both fundamental (e.g., length, mass, time) and compound (e.g., force, energy, charge), with built-in extensibility to accommodate new physical quantities.
4. ***Accuracy***: High-precision unit conversions and computations, particularly when handling data of extreme magnitudes.
5. ***Efficiency***: Optimized implementation to minimize computational overhead for unit checking, especially in large-scale numerical operations.
6. ***Type safety***: Prevention of illegal operations between incompatible units, such as direct addition of time and length.

Furthermore, to function as a unit system supporting HPC capabilities, `BrainUnit` was designed with full compatibility for JAX in mind. Its goals are to: (1) seamlessly integrate JAX’s AD functionality, ensuring that units maintain both the accuracy and efficiency of gradient computations; (2) fully support JAX’s Just-In-Time (JIT) compilation and GPU/TPU acceleration, without diminishing optimization benefits or increasing execution time; (3) efficiently manage JAX’s vectorized and parallelized operations; and (4) minimize any negative impact on computation efficiency, thus preserving high-performance computing outcomes.

The following section elucidates our approach to implementing these design principles.

## 2 Implementation

`BrainUnit`’s innovation centers on three interconnected data structures: `brainunit.Dimension`, `brainunit.Unit`, and `brainunit.Quantity` (Figure 1; Sections 2.1–2.3). These structures are complemented by an associated unit-aware mathematical system (Figure 2; Sections 2.4–2.6).

**Figure 1.**
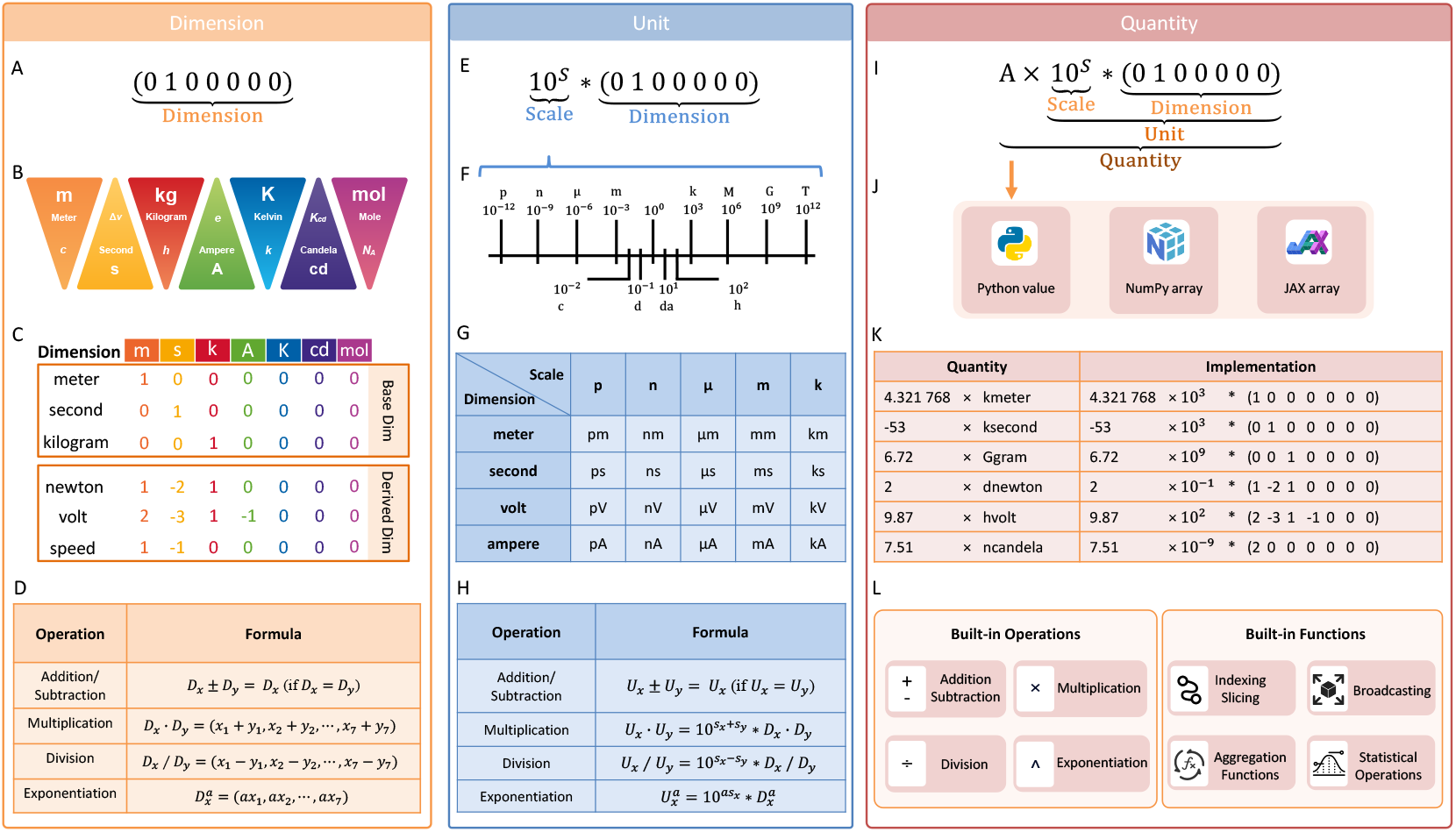
Core Data Structures in BrainUnit for Physical Unit Processing. (A-D) `brainunit. Dimension` enables dimensional analysis and automatic unit conversion based on SI unit rules. (A) The `Dimension` data structure represents the dimensionality of the seven base SI units shown in (B). The SI base units include: meter (`m`) for length, second (`s`) for time, kilogram (`kg`) for mass, ampere (`A`) for electric current, kelvin (`K`) for temperature, candela (`cd`) for luminous intensity, and mole (`mol`) for amount of substance. (C) Examples of `Dimension` instances for both base and derived units. Base dimensions are represented by one-hot vectors, where the position of a “1” corresponds to the relevant SI unit in (B). Derived dimensions are formed by combining these vectors as described in (D). (D) Operational rules for `Dimension` instances, including addition, subtraction, multiplication, division, and exponentiation. (E-H) `brainunit.Unit` provides representations for physical units. (E) The `Unit` data structure is composed of a metric scale and a `Dimension` instance. While the default base factor is 10, custom base factor can be specified. (F) The metric scale varies widely in physical worlds, rangging from pico-to terascale. By adding this scale prefix, `Unit` can represent entities as small as atoms or as large as stars. (G) Examples of `Unit` instances for representing various physical units. (H) Operation rules for `Unit` instances, including addition, subtraction, multiplication, division, and exponentiation. (I-L) `brainunit.Quantity` provides an array-based interface for unitaware numerical computations. (I) The `Quantity` data structure is composed of a numerical value *A* and a `Unit` instance. (J) The numerical value *A* can be implemented as Python numbers, NumPy arrays, or JAX arrays. (K) Examples of `Quantity` instances for representing physical quantities used in scientific computing. (L) Supported mathematical operations for `Quantity` include the basic arithmetic functions (e.g., addition, subtraction, multiplication, division, and exponentiation) and commonly used array functions (e.g., indexing, slicing, broadcasting), following the NumPy array syntax [18].

**Figure 2.**
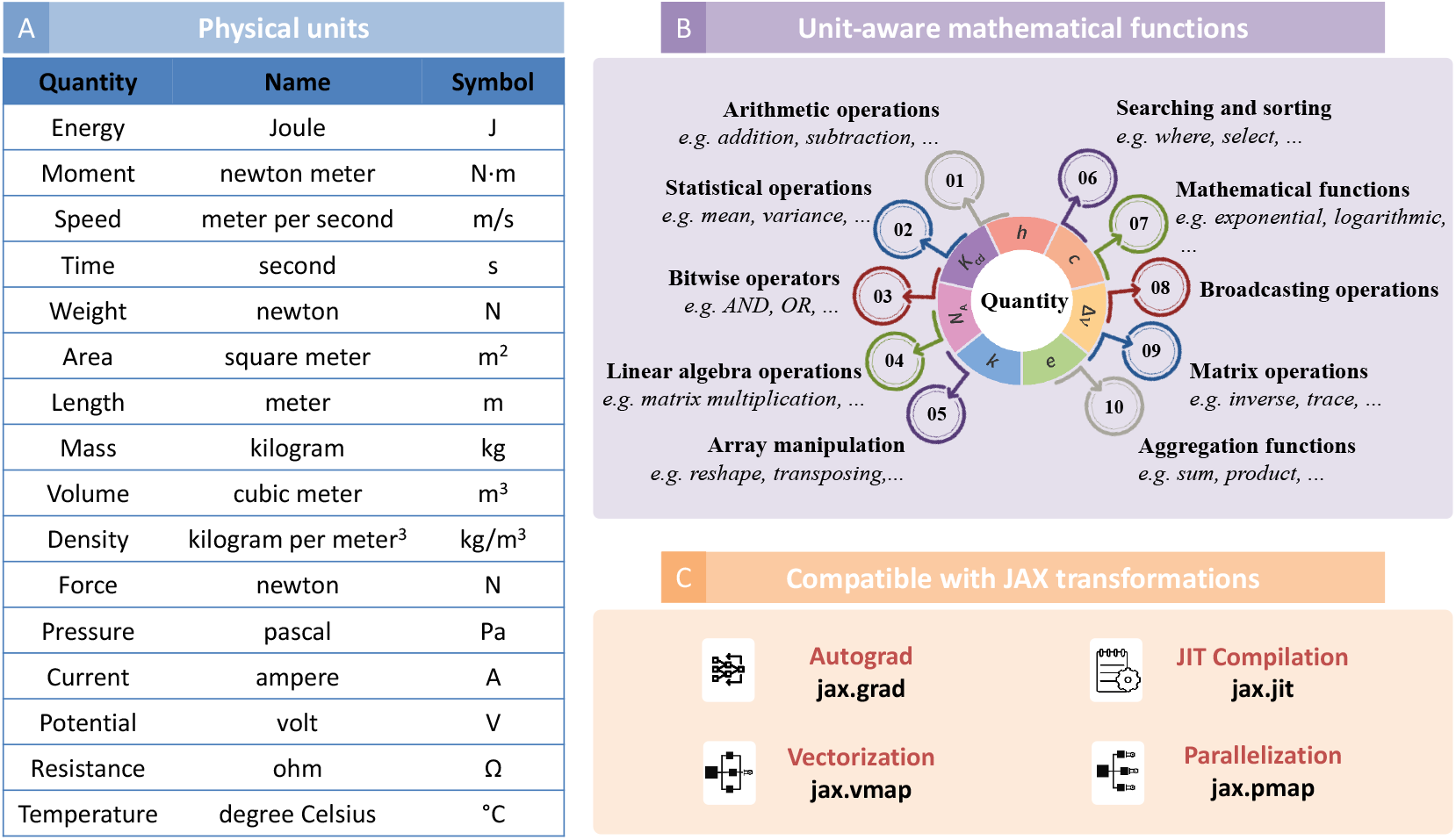
JAX-Compatible Physical Units and Unit-Aware Mathematical Functions in BrainUnit. (A) Categories of physical units supported by `BrainUnit`, spanning over 2,000 commonly used units in fields such as physics, chemistry, astronomy, geoscience, biology, psychology, materials science, and engineering.. (B) Unit-aware mathematical operators in `BrainUnit` follow NumPy’s array programming conventions. Supported operations include but not limited to arithmetic, statistics, bitwise manipulation, linear algebra, array manipulation, searching, sorting, broadcasting, aggregation, and matrix operations. (C) All physical units and mathematical functions in `BrainUnit` are fully compatible with JAX transformations, including `jax.grad` for automatic differentiation, `jax.vmap` for vectorization, `jax.pmap` for parallelization, and `jax.jit` for JIT compilation.

### 2.1 brainunit.Dimension for Automatic Unit Conversion

`Dimension` establishes a rigorous unit conversion system. It is based on the SI Units [4], the most widely adopted standard measurement system globally. This system is founded on seven fundamental physical units: length (`m`), mass (`kg`), time (`s`), electric current (`A`), thermodynamic temperature (`K`), amount of substance (`mol`), and luminous intensity (`cd`) (Figure 1B). In our implementation, `Dimension` is represented as a tuple of seven integers (Figure 1A). Each integer in this tuple corresponds to the dimension of a base unit, where the value indicates the number of dimensions of that particular base unit. Derived units, which measure other physical quantities such as force, electric charge, and speed, are created through mathematical combinations of these fundamental one-hot base vectors (Figure 1C).

The composition rules for `Dimension` follow the SI rule and are illustrated in Figure 1D. These rules govern how dimensions interact under various operations. (1) Addition and subtraction of two `Dimension`s are only defined for those of the same type, otherwise a mismatch error will be raised. The resulting dimension remains unchanged, as these operations do not alter the fundamental units involved: **D**_*x*_ *±* **D**_*y*_ = **D**_*x*_. (2) Multiplication between two `Dimension`s is performed by adding their corresponding elements: **D**_*x*_ · **D**_*y*_ = (*x*_1_ + *y*_1_, *x*_2_ + *y*_2_, …, *x*_7_ + *y*_7_). (3) Division is achieved by subtracting the elements of the denominator from those of the numerator: **D**_*x*_*/***D**_*y*_ = (*x*_1_ *™ y*_1_, *x*_2 *™*_ *y*_2_, …, *x*_7_ *™ y*_7_). (4) Exponentiation with a scalar power *s* is performed by multiplying each element by the scalar: 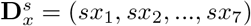 These composition rules ensure that all unit conversions and calculations maintain dimensional consistency.

### 2.2 brainunit.Unit for Physical Unit Representations

Physical phenomena and objects can vary dramatically in size and magnitude. For instance, lengths can range from the subatomic scale of particles to the astronomical scale of galaxies. While the standard metric unit of length is the meter, it is often impractical to use this base unit for all measurements. Real physical units frequently incorporate prefixes that indicate their scale relative to the base unit. These prefixes are based on powers of 10, both positive (10^1^, 10^2^, 10^3^, etc.) and negative (10^−1^, 10^−2^, 10^−3^, etc.), as illustrated in Figure 1F (Table S1 provides a comprehensive list of metric prefixes in the metric system). Therefore, `Unit` is implemented to represent the true physical unit used at the user-level interface. It consists of a metric scale and a `Dimension` instance (Figure 1E). This structure allows for the representation of a wide range of physical quantities across various scales (Figure 1G). Users can dynamically switch to the most appropriate metric prefix, ensuring that quantities are always expressed in the most suitable units for the given context.

The operations between two instances of `Unit` follow five fundamental rules (Figure 1H). (1) Addition and subtraction: These operations are only valid between units with the same dimension. The result maintains the dimension of the operands, but the scale may be adjusted to the most appropriate prefix. (2) Multiplication: When multiplying two units, their dimensions are combined according to the rules of `Dimension` multiplication, and their scales are multiplied: 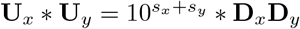 (3) Division: Similar to multiplication, but the dimensions are combined using `Dimension` division, and the scales are divided: 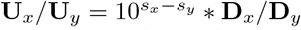. (4) Exponentiation: When a unit is raised to a power, its dimension is exponentiated according to the `Dimension` exponentiation rules, and its scale is raised to the same power: 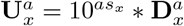. (5) Comparison: Units can only be compared if they have the same dimension. The comparison is performed after converting both units to a common scale. These five rules ensure that all operations on `Unit` instances maintain physical consistency and provide meaningful results across different scales and dimensions.

### 2.3 brainunit.Quantity for Unit-Aware Array Computations

High-performance computing today relies heavily on operations performed on vectors, matrices and higher-dimensional arrays [18]. The `Quantity` structure implements such array-based programming with unit-aware numerical computation. It is implemented as a Python object that integrates a numerical value with a unit (Figure 1I). The numerical value *A* is typically represented as a Python number, NumPy array, or JAX array (Figure 1J), while the unit is instantiated as a `Unit` object. One of the key features of `Quantity` is its flexibility in deployment. The numerical value *A* can be deployed on various computational devices, including CPUs, GPUs, and TPUs, allowing for efficient computation across different hardware architectures. In contrast, the unit component, being a `Unit` instance, operates exclusively on the CPU at compile time, facilitating unit checking without impacting runtime performance. `Quantity` naturally aligns with scientific notation (*x ×* 10^*y*^), where the numerical value *A* corresponds to the mantissa *x*, and the scale of the unit represents the exponent *y*. However, it extends beyond traditional scientific notation by incorporating an additional layer of dimensional analysis (Figure 1K).

Unlike traditional physical unit packages (see Related Works), `Quantity` employs a unique approach to storing quantities. Instead of multiplying the metric scale *s* into the numerical value *A*, it maintains separate representations for the numerical value *A*, metric scale *s*, and dimension. This design choice offers significant advantages, particularly when *A* is represented using low-precision data types like float16, which are common in deep learning applications. By keeping the metric scale separate, `Quantity` can handle very large (e.g., exa-, 10^24^) or very small (e.g., yocto-, 10^−24^) scales without compromising computational accuracy or encountering the limitations typically associated with low-precision arithmetic.

To ensure seamless integration with existing scientific computing workflows, `Quantity` has overloaded nearly all mathematical operators supported by NumPy and JAX arrays (Figure 1L). This comprehensive overloading enables automatic unit processing across a wide range of operations, including basic arithmetic operations (addition, subtraction, multiplication, division), indexing and slicing, broadcasting, aggregation functions (maximum, minimum, sum, mean, product), statistical operations (standard deviation, variance), and many more.

### 2.4 Comprehensive Physical Units

`BrainUnit` offers a comprehensive library of over 2000 commonly used physical units (see `brainunit` `.Unit` for Physical Unit Representations). This extensive collection encompasses units from various domains of physics and engineering, including but not limited to: mass, angle, time, length, pressure, area, volume, speed, temperature, energy, power, and others (details please see Figure 2A and Table S2). By providing such a wide array of pre-defined units, `BrainUnit` significantly reduces the burden on researchers and engineers to manually define and manage units in their calculations, thereby minimizing errors and improving the efficiency of scientific computations. Moreover, the library is designed to be extensible, allowing users to define and add their own custom units if needed, further expanding its applicability to specialized fields or unique experimental setups (see SI A).

### 2.5 Unit-Aware Mathematical System

`BrainUnit` is equipped with a robust unit-aware mathematical framework capable of automatically handling the diverse range of physical quantities it represents (see `brainunit.Quantity` for Unit-Aware Array Computations) when performing complex numerical computations. In the Python scientific computing ecosystem, mathematical array programming has been standardized and popularized by NumPy [18]. Recognizing the widespread adoption and familiarity of this NumPy paradigm, `BrainUnit` has been designed to seamlessly integrate with existing workflows. Particularly, `BrainUnit` provides an extensive set of over 300 commonly used mathematical operators that closely mirror the syntax and behavior of their NumPy counterparts. This design choice allows most Python users to leverage their existing knowledge and immediately start programming in `BrainUnit` using familiar mathematical operator syntax, while benefiting from the added dimension of unit-aware calculations. The rich set of operators provided by `BrainUnit` encompasses a wide range of mathematical and array manipulation functionalities, see illustrations in Figure 2B. Based on them, `BrainUnit` ensures to handle a wide variety of scientific computations while maintaining unit consistency throughout the calculations.

### 2.6 Compatibility with JAX Transformations

JAX has emerged as a high-performance numerical computing library, gaining widespread adoption in fields such as machine learning and scientific computing. Its power lies in a set of composable transformations that enable advanced high-performance computation, including `jax.grad` for autograd, `jax.jit` for JIT compilation, `jax.vmap` for vectorization, and `jax.pmap` for multi-device parallelization. To leverage these powerful transformations, `BrainUnit` has been implemented with full consideration of JAX compatibility (Figure 2C). A key aspect of JAX’s design is its use of the `PyTree` data structure. Any dynamic data within a `PyTree` can be differentiated, compiled, vectorized, or parallelized using JAX transformations. Recognizing this, we have registered `Quantity` as a `PyTree`, in which the numerical value *A* is treated as dynamic data, while the unit is considered static data. This separation offers several significant advantages. (1) By treating only the mantissa *A* as dynamic data, gradient computation is restricted to this numerical value, rather its unit. This approach prevents illegal or nonsensical gradient definitions on metric scales *s*, which are inherently non-differentiable. (2) It allows JIT compilation to trace only the mantissa values. Unit conversion and checking only occur during compilation time rather than runtime after compilation. This design minimizes the overhead of physical unit processing on numerical computation, resulting in maximized performance comparable to raw numerical operations. (3) By treating the unit as static data, `BrainUnit` ensures that unit information is preserved on data throughout JAX transformations. This prevents accidental unit loss or corruption after gradient or JIT computation.

## 3 Evaluations

Brain dynamics modeling creates models span multiple scales, from molecular interactions to wholebrain networks, and are intrinsically linked to empirical data. It is characterized by the use of complex and diverse physical units, making it an ideal testbed for demonstrating `BrainUnit`’s capabilities in handling multi-scale unit conversions and calculations. Therefore, we conducted a series of experiments using a diverse array of brain dynamics models to assess the efficacy of `BrainUnit` in terms of unit consistency, performance overhead, and practical applicability.

### 3.1 Use Case 1: Hodgkin-Huxley Neuron Models

A typical use case for `BrainUnit` is the modeling of detailed biophysical Hodgkin-Huxley-style neuron models. These biological neuron models usually incorporate various physical units, including voltage (e.g., `millivolt`, mV), current (e.g., `nanoamp`, nA), time (e.g., `millisecond`, ms), length (e.g., `micrometre`, mm), concentration (e.g., `mole`, mol/L), etc. These units are frequently utilized across computer simulations and experimental data processing. The introduction of `BrainUnit` is excepted to ensure the correctness and consistency of these units in the computational process, thereby reducing computational errors and improving the model accuracy. To demonstrate this feature, we show how `BrainUnit` can be integrated into `Dendritex`, a package designed to construct detailed multi-compartment neuron models for modeling ion flows, ion channel dynamics, synaptic inputs and membrane potential propagation.

Figure 3A illustrates how `BrainUnit` aids `Dendritex` in constructing a biophysical complex multichannel neuron model of the thalamus reticular nucleus (TRN) [5, 6]. This neuron model includes five active ionic currents: a spike generating fast sodium current (*I*_Na_), a delayed rectifier potassium current (*I*_DR_), a low-threshold T-type Ca^2+^ current (*I*_Ca*/*T_), a Ca^2+^-dependent potassium current (*I*_AHP_) and a Ca^2+^-activated nonselective cation current (*I*_CAN_). The integration of `BrainUnit` not only makes the code more readable and intuitive but also ensures the direct interoperation with empirical data, enhancing both theoretical and practical research capabilities.

**Figure 3.**
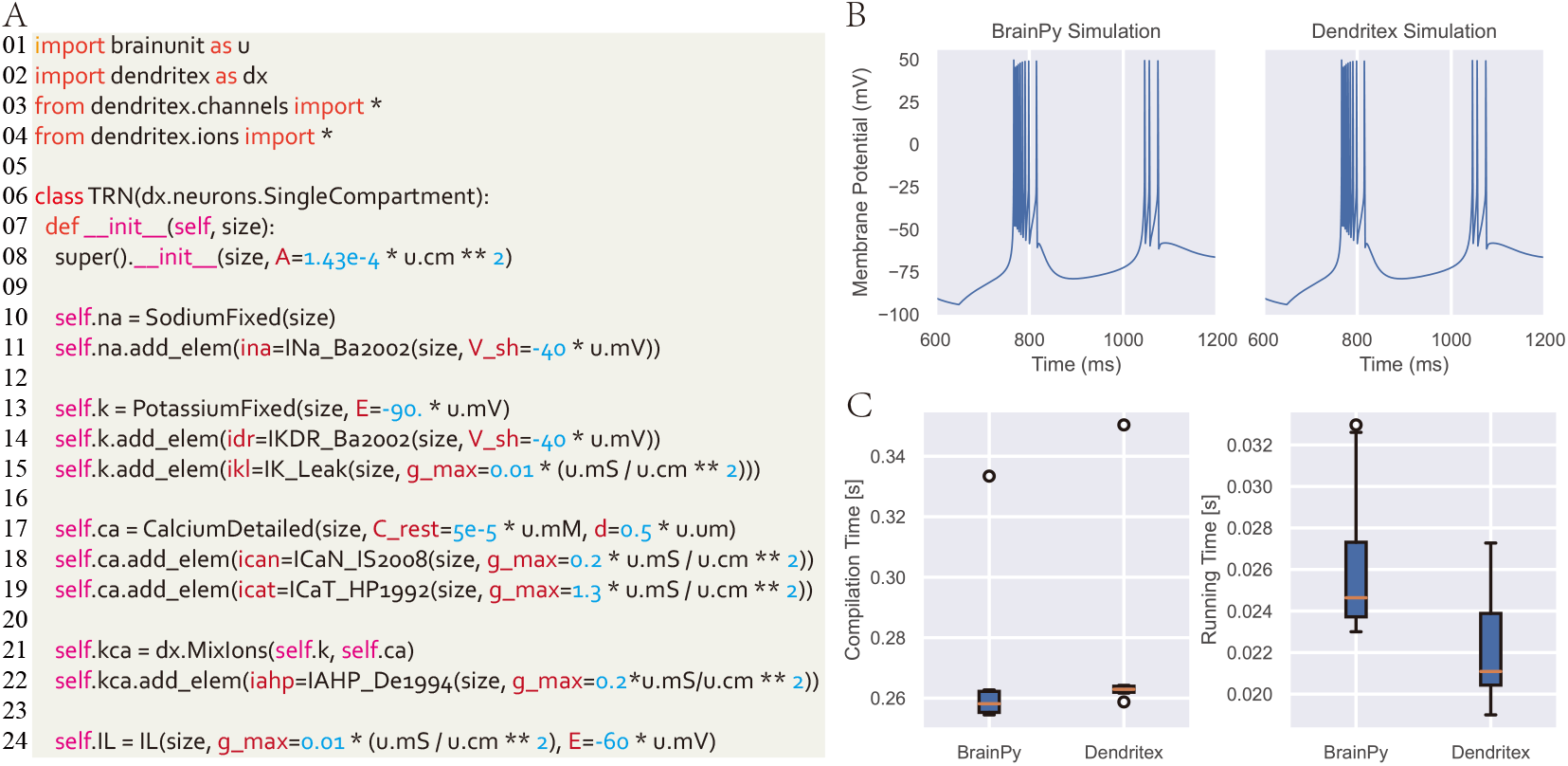
Modeling Biologically Detailed Hodgkin-Huxley Neuron Models with BrainUnit. (A) Integration of physical units in `Dendritex` enables precise modeling of biologically detailed Hodgkin-Huxley neuron models. By embedding unit-aware operations directly into the neuron simulation, `Dendritex` ensures accurate and meaningful calculations, reducing the risk of unit-related errors in complex biological simulations. (B) Comparison of simulation results between `BrainPy` (without physical unit tracking) and `Dendritex` (with unit-aware functionality). The introduction of physical units in `Dendritex` does not compromise simulation accuracy but adds clarity and error prevention by automatically handling unit conversions and ensuring dimensional consistency throughout the computation. (C) Performance comparison between `BrainPy` and `Dendritex`. The left panel shows the compilation time, while the right panel highlights the simulation time. While the inclusion of physical units in `Dendritex` introduces a slight overhead in both compilation and simulation, the benefits of improved accuracy and unit-safety far outweigh the minimal performance cost, making it highly suitable for biologically detailed neuron modeling.

We conducted a comparative analysis of simulation results obtained from `Dendritex` and `BrainPy`, a differentiable brain simulator [39, 40] that lacks automated unit processing. Both simulations yielded identical voltage traces, as illustrated in Figure 3B, demonstrating the accuracy and reliability of `Dendritex`’s unit-aware computations. While `Dendritex` exhibited a slightly longer compilation time due to the additional processing required for unit conversion, its post-compilation performance was comparable and even superior to that of `BrainPy` (Figure 3C).

### 3.2 Use Case 2: Multiscale Brain Models

Modern scientific computing usually does not focus on a single scale, but integrate data across multiple scales. For example, brain dynamics modeling needs to integrate the behavior of models from molecular to cellular, network, and up to the level of entire nervous networks. This multiscale integration requires dealing with units at different levels, which `BrainUnit` can efficiently manage and transform, allowing for more accurate and reliable modeling from the microscopic to the macroscopic.

To demonstrate this feature, we show how `BrainUnit` can be integrated into `BrainPy` to help build multi-scale brain dynamics networks.

Leveraging the connectome data of the macaque brain [27] (Figure 4A), we developed a large-scale spiking network model that captures the hierarchical organization of the brain. This model spans various scales, from synapses and neuronal populations to networks and circuits (Figure 4B). It includes 29 distinct brain regions, each modeled as an excitatory-inhibitory (EI) balanced network consisting of 3200 excitatory neurons and 800 inhibitory neurons. These neurons are connected through current-based exponential synapses, allowing for biologically realistic neuronal interactions. Physical units are embedded at every level of the model to enhance its accuracy and relevance (Figure 4B). For example, at the neuronal and synaptic levels, the `BrainUnit` framework enables precise coding of membrane potentials and time constants. At the network level, it captures detailed synaptic connectivity and network dynamics, while at the circuit level, it accounts for essential circuit parameters such as areal distances, synaptic conduction velocities, and delays, all standardized through `BrainUnit`.

**Figure 4.**
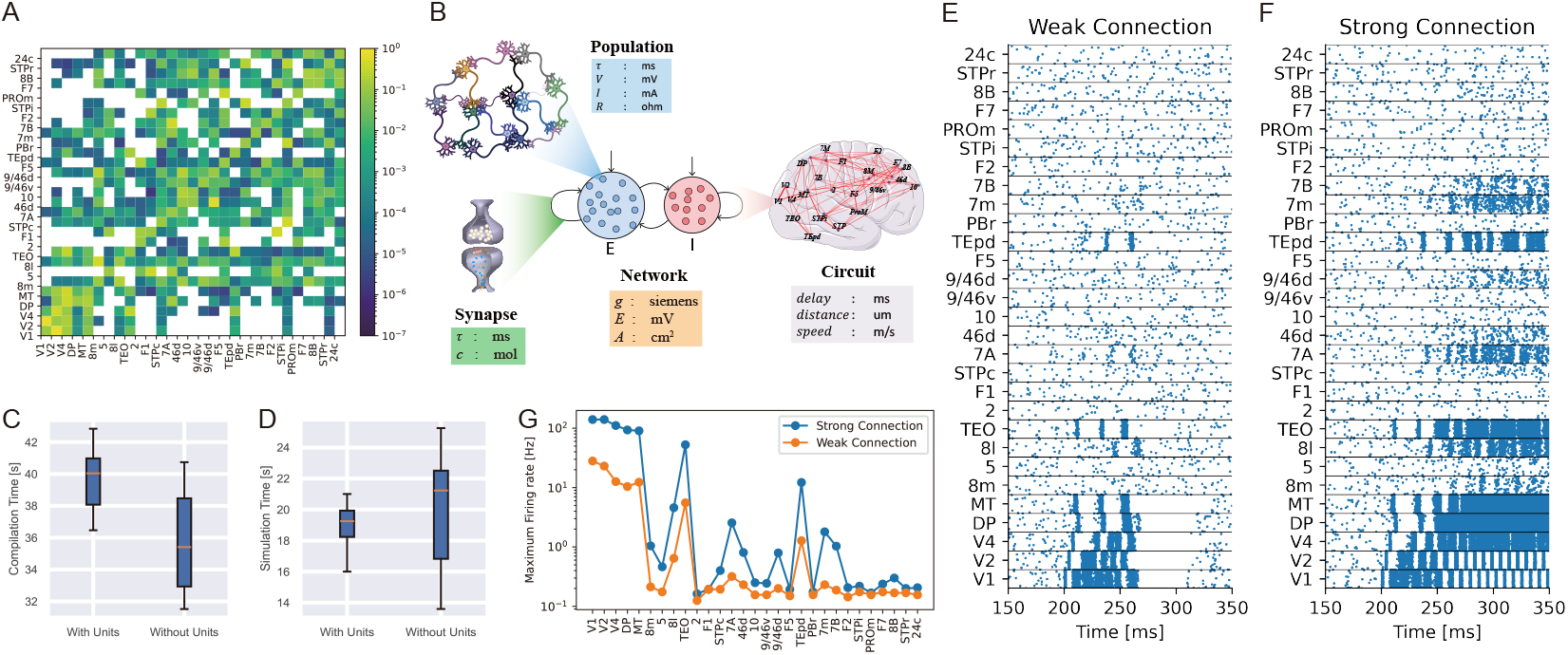
Multiscale Brain Modeling with BrainUnit. (A) Connectivity probabilities among 29 brain areas, as measured in Markov et al. [27]. Each cell represents the probability of connection between a presynaptic brain area (row) and a postsynaptic brain area (column). White grid cells indicate zero connectivity. (B) Integration of physical units into multiscale brain models. At the synaptic level, physical units define parameters like synaptic time constants and neurotransmitter concentrations. At the population level, they describe membrane potentials and synaptic currents. At the network level, physical units quantify synaptic weights and network area. At the circuit level, they capture metrics such as synaptic delays, inter-area distances, and transmission speeds. (C)Comparison of compilation times for multiscale brain models with and without physical units. (D)Comparison of simulation times for multiscale brain models with and without physical units. (E)Raster plot of neural activity across 29 brain areas when inter-area connection strength is weak. The network responds to a 50 ms pulse input to V1, and the activity propagates along the cortical hierarchy. Brain areas in the ventral stream show strong responses. (F) Raster plot of neural activity across 29 brain areas when inter-area connection strength is strong. Similar to (B). The activity propagates more extensively, reaching higher-order cortical areas, including the prefrontal cortex. (G) Peak firing rates across 29 brain areas, comparing weak (orange) and strong (blue) inter-area connection strengths. The propagation of neural activity from V1 is more pronounced and widespread in the strong connection scenario.

To evaluate the model, we first compared compilation and simulation times with and without unitaware modeling. Our findings showed that incorporating physical units introduced a minor compilation overhead, though it was not significant (Figure 4C). Notably, the simulation speed remained comparable with and without units (Figure 4D). We then tested simulation accuracy by applying a brief (50 ms) pulse input to the early visual cortex (V1). The resulting raster plots revealed nearly identical neuronal activity in both scenarios (these plots are not shown as they are visually indistinguishable). When inter-area excitatory coupling was weak (*µ*_EE_ < 0.08 mS), activation was limited to a subset of the prefrontal cortex (PFC), particularly the frontal eye fields (area 8l) (Figure 4E). Under these conditions, signal propagation occurred primarily along the ventral visual stream (V1, V2, V4, TEO, and TEpd), driven by stronger anatomical projections within these regions. However, when the inter-area excitatory coupling was increased (Figure 4F), we observed heightened activity in higher brain areas associated with advanced cognitive functions [17]. These regions included the dorsolateral prefrontal cortex (dLPFC; areas 46d, 9/46d), the frontopolar cortex (area 10), the parietal cortex (areas 7A, 7B, and 7m) in the dorsal stream, and the frontal eye fields (areas 8l, 8m). This was reflected in the peak firing rates observed across regions (Figure 4G). These results demonstrate that incorporating physical units into multiscale brain modeling not only enhances the biological realism and interpretability of simulations but also ensures accurate representation of emergent brain functions, from local circuit dynamics to large-scale activity patterns across multiple regions.

### 3.3 Use Case 3: Neuronal Activity Fitting

Theoretical models in scientific computing often need to use experimental data to validate and adjust their parameters. Experimental data contains specific physical units, which theoretical models need to handle accurately. The application of `BrainUnit` simplifies this process by ensuring consistency and accuracy of the data units, thus improving the predictive power and reliability of the models. To demonstrate this feature, we show how `BrainUnit` can be integrated into traditional optimization methods such as differential evolution [34] for fitting Hodgkin-Huxley models to experimentally collected voltage clamp data.

The experimental data comes from the in *vitro* intracellular recording of a cortical pyramidal cell. Five different step currents were injected into the cell (three inputs are shown in bottom panels of Figure 5A-C), and the membrane potentials were recorded simultaneously. We constructed a Hodgkin-Huxley neuron model, and we fitted the parameters of Sodium, Potassium, and leakage channel conductance, and the membrane capacitance. Prior to the fitting process, the user is required to define the bounds for each fitting parameter (Figure 5D), after which the optimization process is fully automated. We compared the fitting speed and results with `brian2modelfitting`, a toolbox for data driven optimization in Brian2 simulator [33]. We found that the fitting with `Dendritex` (middle panels in Figure 5A-C) achieved better accuracy than the fitting in `brian2modelfitting` (top panels in Figure 5A-C). Quantitative comparison revealed fitting with `Dendritex` requires more time than `brian2modelfitting` (Figure 5E), while `Dendritex` achieves overall much lower loss function, indicating the better fitting results are achieved (Figure 5F). These results highlight the effectiveness of `Dendritex` for achieving higher-precision parameter fitting in neuron models, despite the trade-off in computational speed.

**Figure 5.**
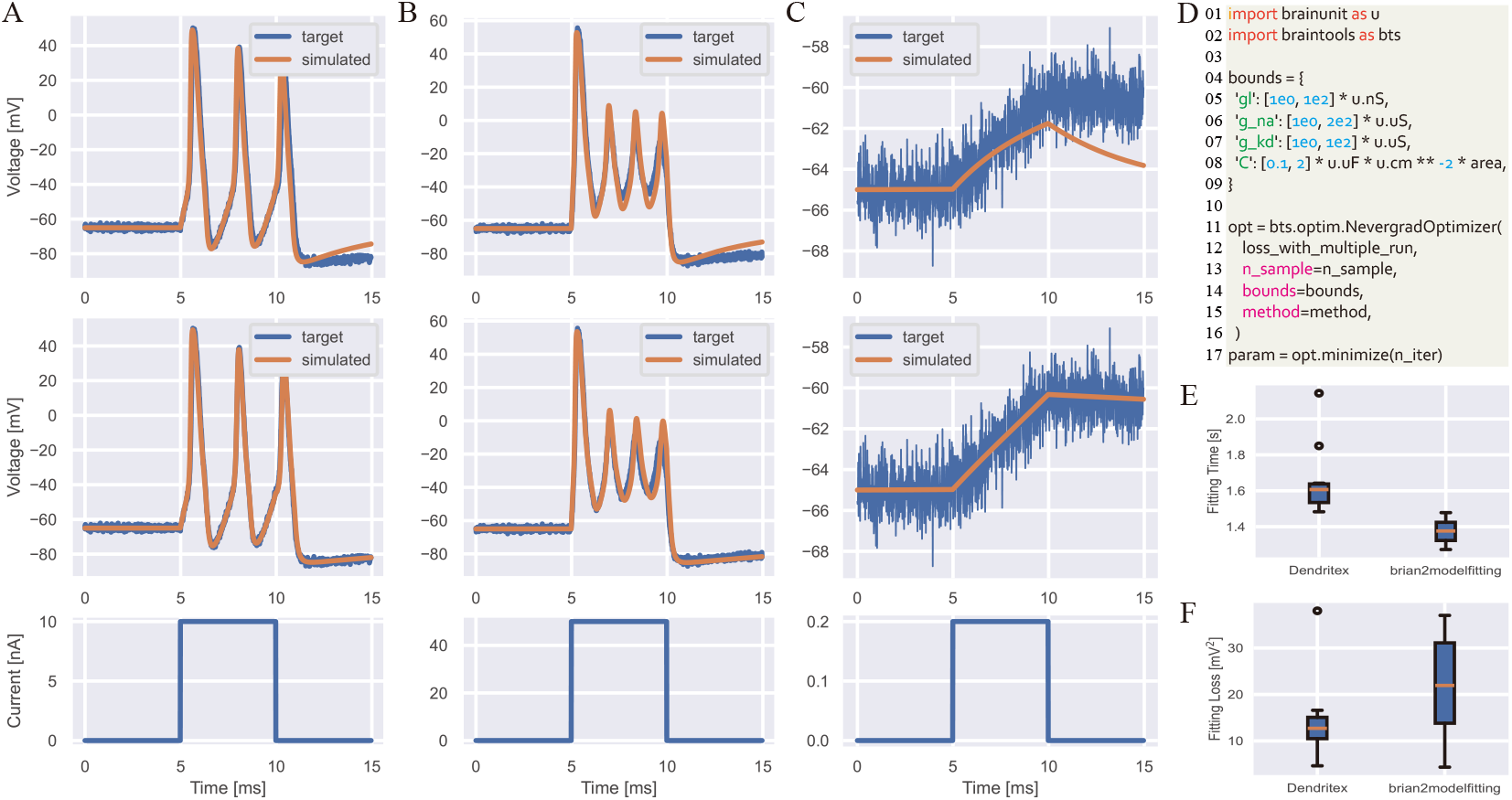
Fitting Single-Neuron Activity Using the Hodgkin-Huxley Neuron Model within BrainUnit. (A-C) Comparison of fitting results between `brian2modelfitting` and `BrainUnit`. The top row displays the fitting results from `brian2modelfitting`. The middle row shows the fitting results from `BrainUnit`. The bottom row represents the input current applied to the cortical pyramidal cell. (D) Integration of physical units into traditional non-differentiable optimization methods, such as differential evolution [34], within `BrainUnit`. (E) Comparison of fitting times between `brian2modelfitting` and `BrainUnit`. (F) Comparison of fitting losses between `brian2modelfitting` and `BrainUnit`.

### 3.4 Use Case 4: Cognitive Task Training

In scientific computing, we heavily utilize the automatic differentiation capabilities in HPC libraries to address higher-order optimization problems. For instance, in brain dynamics modeling, neural networks are commonly trained to perform cognitive tasks using gradient-based optimization methods [31]. Therefore, it is essential that an AI-compatible unit system supports classical AD techniques.

`BrainUnit` seamlessly integrates with both forward and backward AD methods supported in JAX [10], specifically the vector-Jacobian product (VJP) and Jacobian-vector product (JVP). During forward computation, the unit system ensures consistency among data units as demonstrated in above sections (Figure 6A). However, during the AD pass, the unit checking will be closed and do not happen among gradients. No matter in VJP (Figure 6B) or JVP (Figure 6C) gradient passes, the computed gradients will be associated with the same unit as the original data. This feature that treating the unit as the static data and not flowing through gradient computation are very useful to be directly compatible with traditional AD optimizers.

**Figure 6.**
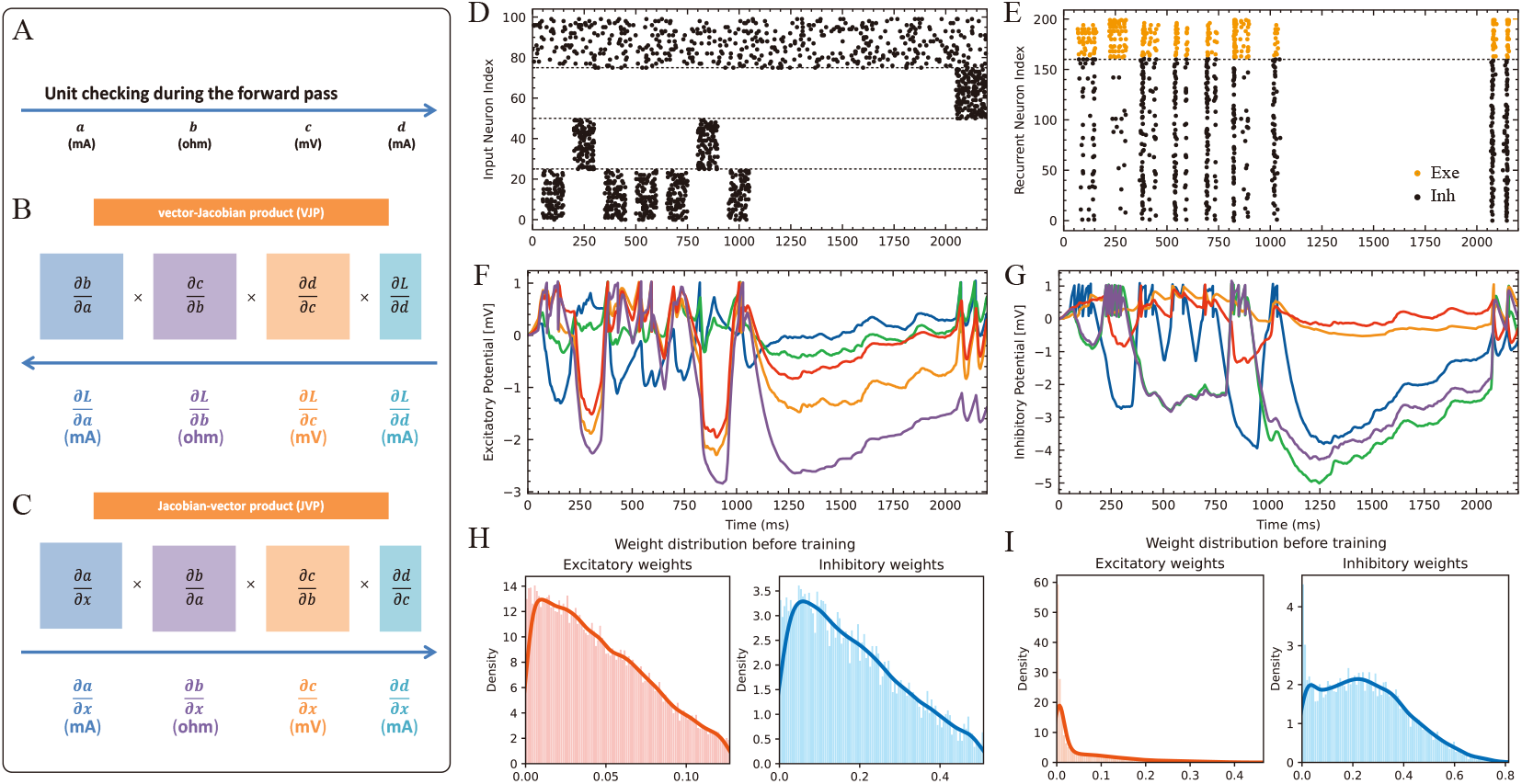
Training Cognitive Tasks on a Biological EI Network Powered by BrainUnit using Differentiable Optimization. (A) In a gradient optimization task, `BrainUnit` ensures physical unit consistency during the forward pass of model computation. During the gradient computation pass, whether using backward-mode (B) or forward-mode (C) automatic differentiation, unit checking is disabled. However, the computed gradients retain the physical units associated with the data from the forward pass. (B) Physical units are linked to the gradients computed using the backward-mode vector-Jacobian product rule in automatic differentiation. (C) Physical units are linked to the gradients computed using the forward-mode Jacobian-vector product rule. (D) Example of an input spike train used in an evidence accumulation task. (E) Example spike train of excitatory and inhibitory recurrent units in response to one trial of the task shown in (D). (F) Five examples of membrane potential traces from excitatory recurrent units during task processing. (G) Five examples of membrane potential traces from inhibitory recurrent units during task processing. (H) Distribution of excitatory and inhibitory synaptic weights in the network before training. (I) Distribution of excitatory and inhibitory synaptic weights in the network after training.

To access the efficacy of `BrainUnit` within such gradient-based frameworks, we integrated it into the online learning framework `BrainScale` [37], testing its performance and accuracy when training biologically-inspired excitatory-inhibitory (EI) networks for cognitive tasks [38]. This model incorporates generalized integrate-and-fire (GIF) neurons [22] to govern spiking dynamics, while synaptic interactions are simulated using the conductance-based exponential model. We trained this unit-enhanced EI network on evidence accumulation tasks [29] (Figure 6D) using the `BrainScale` platform [37]. Our analysis revealed that incorporating physical units did not alter the fundamental network dynamics post-training. During the evidence accumulation phase, we observed distinct activation patterns: inhibitory neurons showed pronounced responses following stimulus presentation, while excitatory neurons maintained comparatively lower firing rates (Figure 6E). This behavior aligns with the expected dynamics of balanced EI networks in cognitive tasks. Examination of neuronal membrane potentials (Figure 6F-G) highlighted a key advantage of our conductance-based modeling. Unlike current-based synapse models prevalent in deep learning applications, this model naturally constrained membrane potentials between excitatory and inhibitory reversal potentials, eliminating the need for artificial voltage regularization techniques. We further investigated synaptic weight distributions before and after training (Figure 6H-I). Initial weights for both excitatory and inhibitory connections were drawn from the absolute values of a normal distribution (Figure 6H). Post-training, the weight distributions evolved to closely mimic biological observations: excitatory weights approximated the tail of a Gaussian distribution [7], while inhibitory weights exhibited a log-normal distribution [26, 12]. Overall, these results demonstrate that integrating physical units into gradient-based modeling maintains computational efficiency while preserving biological realism, offering a promising approach for more accurate and interpretable neural simulations.

## 4 Discussion

In conclusion, we introduced `BrainUnit`, a unit system designed to seamlessly integrate physical units into high-performance, AI-driven scientific computing, with a particular emphasis on compatibility with JAX. `BrainUnit` addresses a critical gap in current AI libraries: the lack of native support for physical units, which is vital for rigorous scientific computations. By offering a comprehensive library of over 2000 physical units and an extensive set of unit-aware mathematical operators, `BrainUnit` provides a robust tool for unit-aware scientific computing. Its full integration with JAX transformations allows users to utilize advanced features such as automatic differentiation, JIT compilation, and parallelization, while maintaining unit consistency. Through a series of use cases in brain dynamics modeling, we demonstrated `BrainUnit`’s ability to automate unit processing with minimal performance overhead and its practical applicability in real-world scenarios.

`BrainUnit` represents a significant advancement in the realm of AI-driven scientific computing. While traditional AI libraries excel in high-performance computing with abstract numbers, `BrainUnit` introduces a rigorous physical unit processing engine. By combining the advantage of physical unit handling and high-performance numerical computing, `BrainUnit` enables accurate, reliable, maintainable, interpretable, and rigorous scientific computing. This synergy can be expressed as:

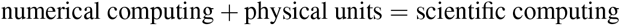

`BrainUnit` offers several key benefits. First, physical units mitigate the risk of unit mismatch, ensuring that results conform to physical laws and improving the correctness of models. Furthermore, explicit unit labeling enhances code readability and maintainability, making it easier for developers to understand variable meanings, particularly in large-scale or interdisciplinary projects. The use of standardized units fosters code portability, allowing libraries from different domains to adopt a consistent unit system, thereby promoting scientific collaboration. Additionally, automatic unit conversion alleviates the burden on developers, minimizing errors from manual conversions and streamlining debugging processes. This not only improves the precision of computations but also accelerates project timelines. Moreover, integrating physical units increases the reliability of simulation outcomes, ensuring that computational models in domains like climate science or aerospace meet real-world physical requirements. Finally, by promoting standardization, `BrainUnit` enhances interoperability between HPC libraries, encouraging broader interdisciplinary collaboration.

`BrainUnit` is specifically designed to integrate with the JAX library [10], although it is easy to extend it to support other AI frameworks. JAX has gained significant popularity in scientific computing due to its rapidly growing ecosystem across multiple domains. Examples include `BrainPy` [39, 40], a differentiable brain simulator, `Brax` [16], a differentiable physics engine, `JAX-COSMO` [13] for cosmological computations, and `JAX-MD` [32] for molecular dynamics simulations. Within this context, the emergence of `BrainUnit` marks a pivotal milestone. It seamlessly integrates the concept of physical units into JAX’s abstract computational framework, paving the way for unit-aware scientific computing in JAX. This advancement will greatly enhance the precision and reliability of JAX computational models across various scientific domains.

Despite the significant benefits `BrainUnit` brings to unit management in AI-driven scientific computing, certain limitations remain that future work could address. Although our evaluations showed minimal performance overhead across different scales of models, there is still a slight compilation overhead associated with unit processing. Future optimizations could further reduce this overhead to enhance performance. While we focused on evaluations based on brain dynamics modeling in this paper, further testing is necessary to validate and optimize `BrainUnit`’s performance across other scientific fields. Additionally, adopting `BrainUnit` in large, pre-existing projects may require substantial refactoring, presenting a potential barrier to widespread adoption. Developing tools to facilitate this transition could significantly lower the adoption barrier. Finally, while `BrainUnit` includes a wide array of physical units, certain specialized or emerging units from niche scientific fields may not yet be supported, highlighting an area for future expansion.

## 5 Methods

### 5.1 Hodgkin-Huxley Models of TRN Neurons

Our single-compartment model of a thalamic reticular nucleus (TRN) cell employed a Hodgkin-Huxley-type framework to model the ionic currents [19]. The membrane potential, *V*, evolved according to the following differential equation [24]:

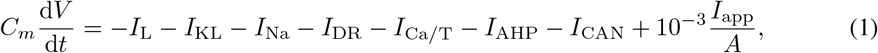

where *C*_*m*_ = 1.0 *µ*F*/*cm^2^ is the membrane capacitance, *I*_L_ is the leakage current, *I*_KL_ is the potassium leakage current, *I*_Na_ is the spike generating fast sodium current, *I*_DR_ is the delayed rectifier potassium current, *I*_Ca*/*T_ is a regular low-threshold T-type Ca^2+^ current, *I*_AHP_ is the Ca^2+^-dependent potassium current, *I*_CAN_ is the Ca^2+^-activated nonselective cation current, *I*_app_ is the externally applied current injection (in nA), and the total membrane area *A* = 1.43 *×* 10^−4^ cm^2^ was used to normalize the externally applied current [15]. Each ionic current was modeled using voltage-dependent activation and inactivation variables, consistent with Hodgkin-Huxley formalism.

The sodium current *I*_Na_ is computed by

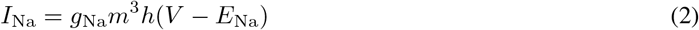

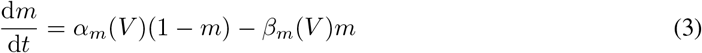

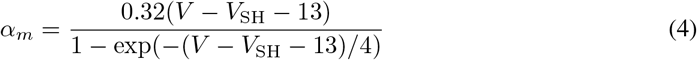

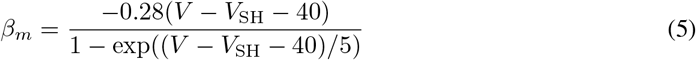

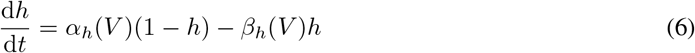

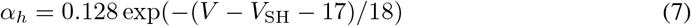

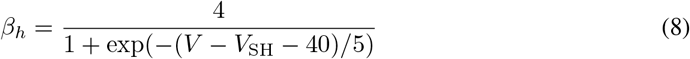

where the maximal conductance *g*_Na_ = 90 mS*/*cm^2^, and the reversal potential *E*_Na_ = 50 mV. *m* is the activation variable, *h* is the inactivation variable. The spike adjusting threshold *V*_SH_ = −40 mV.

The delayed rectifier potassium current *I*_*K*_ is computed by

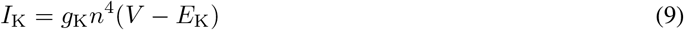

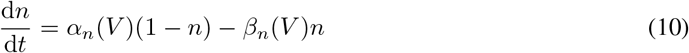

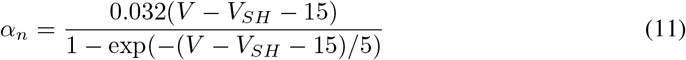

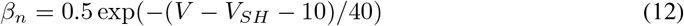

where *V*_SH_ = − 40 mV, the reversal potential *E*_K_ = − 90 mV, and the maximal conductance *g*_K_ = 10 mS*/*cm^2^.

The Ca^2+^-dependent potassium current *I*_AHP_ is computed by:

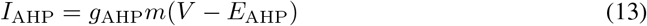

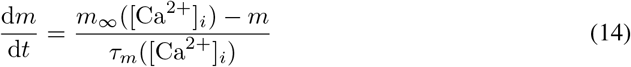

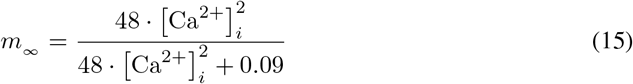

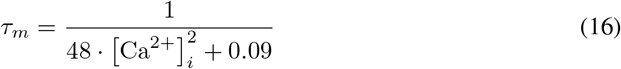

where the reversal potential *E*_AHP_ = − 43 mV, the maximal conductance *g*_AHP_ = 0.2 mS*/*cm^2^, [Ca]_*i*_ is the concentration of the intra-cellular calcium, *E*_Ca_ is the calcium reversal potential.

The Ca^2+^-activated nonselective cation current *I*_CAN_ is computed by:

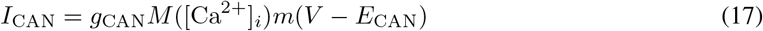

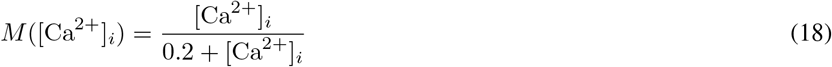

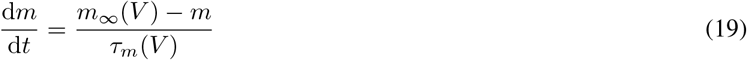

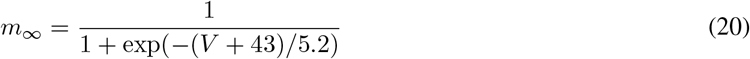

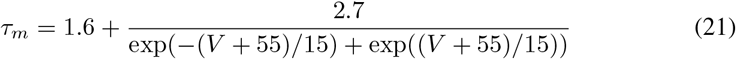

where *M* ([Ca]_*i*_) is a Michaelis-Menten function, the reversal potential *E*_CAN_ = 10 mV, and the maximal conductance *g*_AHP_ = 0.2 mS*/*cm^2^.

The low-threshold T-type Ca^2+^ current *I*_Ca*/*T_ is computed by

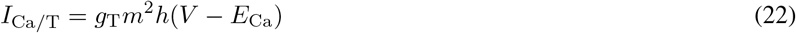

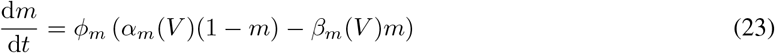

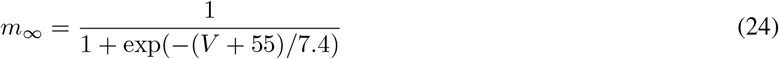

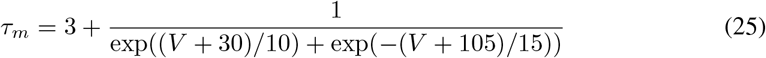

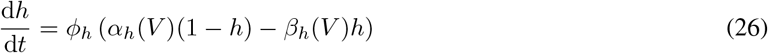

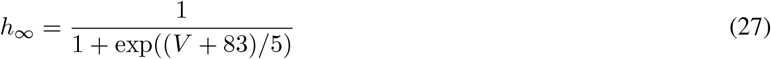

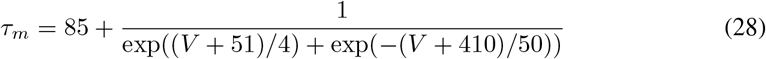

where the maximal conductance *g*_T_ = 1.3 mS*/*cm^2^, *ϕ*_*m*_ = 6.9, and *ϕ*_*h*_ = 3.7. The potassium leaky current *I*_KL_ is computed by

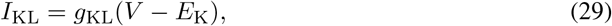

where we set the potassium leakage channel conductance *g*_KL_ = 0.01 mS*/*cm^2^. The leakage current *I*_L_ is computed by

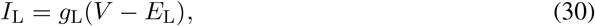

where we used the leakage channel conductance *g*_L_ = 0.01 mS*/*cm^2^ and the leakage reversal potential *E*_L_ = −60 mV.

The calcium reversal potential (*E*_Ca_) was dynamically calculated using the Nernst equation:

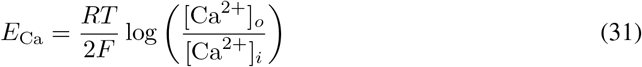

where *R* is the universal gas constant, *T* is the absolute temperature, *F* is the Faraday constant, and [Ca^2+^]_*o*_ and [Ca^2+^]_*i*_ represent the extracellular and intracellular calcium concentrations, respectively. We set [Ca^2+^]_*o*_ = 2 mM.

The intracellular calcium concentration, [Ca^2+^]_*i*_, was governed by a first-order differential equation:

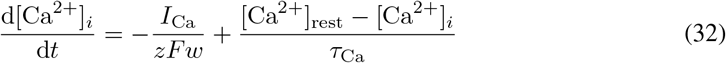

In this equation, *I*_Ca_ = *I*_Ca*/*T_ + *I*_AHP_ + *I*_CAN_ represents the total calcium current, *w* is the effective thickness of the perimembrane shell where calcium influences membrane properties (set to 0.5 *µ*m), *z* = 2 is the valence of calcium ions (Ca^2+^), and *τ*_Ca_ = 100 ms is the time constant for calcium removal. The term [Ca^2+^]_rest_ denotes the resting intracellular calcium concentration, fixed at 0.05 *µ*M. The Faraday constant *F* ensures proper scaling of the ionic flux in relation to the charge carried by Ca^2+^ ions.

The external input current, *I*_app_, used in Figure 3B, follows a step function. For the first 500 ms, *I*_app_ is set to 0 nA. This is followed by a 150 ms period where *I*_app_ is reduced to − 0.05 nA, providing a brief hyperpolarizing input. For the remaining 1000 ms, *I*_app_ returns to 0 nA.

### 5.2 Multi-scale Spiking Neural Network

We developed a spiking network model inspired by the framework of Joglekar et al. [21], incorporating 29 distinct brain areas. The model simulates a large-scale network of leaky integrate-and-fire (LIF) neurons, with both local and long-range connectivity based on the work of Markov et al. [27]. Each brain area is modeled as a population of 4,000 neurons, with a biologically realistic ratio of 3,200 excitatory neurons (80%) and 800 inhibitory neurons (20%). Unlike the delta synapses used in Joglekar et al. [21], synaptic projections in our model follow exponential dynamics. Specifically, synaptic interactions between neurons are governed by the gradual rise and exponential decay of synaptic conductances rather than an instantaneous change. This modification allows for a more biologically realistic representation of synaptic activity over time.

The membrane potential dynamics of each neuron are described by the following equation:

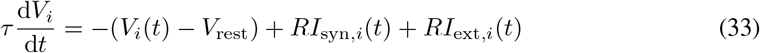

where *V*_*i*_(*t*) is the membrane potential of neuron *i* at time *t, τ* = 20 ms is the membrane time constant, and *R* = 1 Ω is the membrane resistance. The resting membrane potential is *V*_rest_ = − 60.0 mV, and the neuron fires a spike when *V*_*i*_(*t*) exceeds the spike threshold *V*_th_ = − 50.0 mV. Upon spiking, the membrane potential is reset to *V*_reset_ = − 60.0 mV, and the neuron undergoes a refractory period of 5 ms during which it cannot fire again. The term *I*_syn,*i*_(*t*) represents the total synaptic input current, while *I*_ext,*i*_(*t*) denotes any external input current.

The synaptic inputs are modeled using current-based synapses, under the assumption that the membrane potential fluctuates around the resting potential. The synaptic current for neuron *i* is given by:

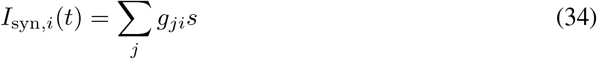

where *g*_*ji*_ is the synaptic conductance from presynaptic neuron *j* to postsynaptic neuron *i*, and *s* = 1 mV is a scaling factor.

The evolution of the synaptic conductance is described by the following equation:

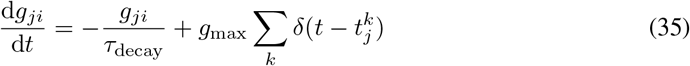

Here, 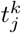 represents the *k*-th spike time of the presynaptic neuron *j*. Whenever a spike occurs in neuron *j*, the synaptic conductance *g*_*ji*_ is increased by a fixed amount, *g*_max_. This conductance then decays exponentially with a time constant *τ*_decay_, which is set to 5 ms for excitatory synapses and 10 ms for inhibitory synapses.

The connectivity structure between neurons follows biologically observed patterns. Intra-areal (within a brain area) connectivity is established based on the excitatory-inhibitory (EI) balance model [11], with a connection probability of 0.1 between neurons within the same area. Inter-areal (between different brain areas) connectivity is determined using the experimental data provided by Markov et al. [27], which quantifies connectivity as a weight index known as the extrinsic fraction of labeled neurons (FLN). This weight index reflects the strength of long-range connections between different cortical areas (Figure 4A).

It is important to note that in Joglekar et al. [21], the connection probability between brain areas was set to be the same as the connection probability within each brain area. In their model, the experimental connectivity data from Markov et al. [27] was used to normalize the connection strength between brain areas, rather than directly determining connectivity patterns. In contrast, our model more closely follows biological measurements by using the experimental connectivity data to explicitly establish the connection matrix between the 29 brain areas. We think this captures the realistic long-range connections, resulting in a more anatomically grounded network structure.

Furthermore, the lengths of projections between brain areas are adapted from the experimental wiring distance matrix of Markov et al. [27], which is based on anatomical measurements of fiber distances. To account for the conduction delay between areas, we incorporate distance-dependent inter-areal synaptic delays, assuming a conduction velocity of 3.5 m/sec [35]. This results in timing delays in signal transmission across the network. Within a brain area, there is no time delay for synaptic transmission.

Within each brain area, the connection strengths *g*_max_ between excitatory (E) and inhibitory (I) populations are specified as follows: the excitatory-to-excitatory (E-to-E) connection strength is *g*_max,EE_ = 0.10108301 S, while the excitatory-to-inhibitory (E-to-I) connection strength is *g*_max,IE_ = 0.60604239 S. The inhibitory-to-excitatory (I-to-E) connection strength is set to *g*_max,EI_ = −0.638 S, and the inhibitory-to-inhibitory (I-to-I) connection strength is *g*_max,II_ = −0.33540355 S.

For inter-areal connections between different brain areas, the excitatory-to-excitatory (E-to-E) connection strength varies depending on the simulation. In the case of a weak connection, the E-to-E connection strength is 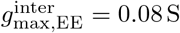 (Figure 4E), while for strong inter-areal connections, the E-to-E strength increases to 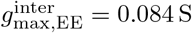 (Figure 4F).

### 5.3 Fitting the Cortical Pyramidal Cell Voltage with the Hodgkin-Huxley Neuron Model

To simulate the electrical behavior of cortical pyramidal cells (the voltage data was adapted from `brian2modelfitting`), we used the Hodgkin-Huxley neuron model [19] as a foundation. The Hodgkin-Huxley model, originally developed to describe the action potential dynamics in squid giant axons, is well-suited for capturing the voltage dynamics of neurons with voltage-gated ion channels. By adjusting the ion channel conductance and membrane capacitance, we can fit the voltage traces recorded from cortical pyramidal cells, allowing for a more accurate representation of their electrophysiological properties.

Cortical pyramidal cells are principal excitatory neurons in the cortex, and their voltage dynamics are shaped by a complex interplay of ion channels, including sodium (Na^+^), potassium (K^+^), and leak channels. The membrane potential *V* of a cortical pyramidal cell is described by a modified Hodgkin-Huxley equation [19] that incorporates these key ionic currents:

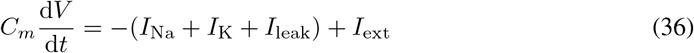

where *C*_*m*_ is the membrane capacitance, *I*_Na_ and *I*_K_ are the sodium and potassium ionic currents, respectively, *I*_leak_ represents the leak current, and *I*_ext_ is the external current applied to the neuron.

Each ionic current is defined by conductance and gating variables that depend on voltage, which determine the opening and closing of ion channels. The sodium current is modeled as Eqs. 2 -8. The potassium current is modeled as Eqs. 9-12. The only difference is the spike adjusting threshold *V*_SH_ is set -63 mV. The leaky current is computed the same as Eq. 30.

The model parameters, including the conductances *g*_Na_, *g*_K_, *g*_L_, and membrane capacitance *C*_*m*_, are fine-tuned by fitting the voltage traces obtained from experimental recordings of pyramidal neurons. To fit the model to the recorded voltage data from cortical pyramidal cells, we use traditional differential evolution [34] to minimize the mean square difference between the simulated voltage and experimental traces.

### 5.4 Conductance-based EI Network for Cognitive Task Training

We developed a biologically inspired conductance-based excitatory-inhibitory (EI) neural network model [37], structured to solve an evidence accumulation task [29]. This architecture is grounded in principles of neuronal dynamics, particularly mimicking the excitation and inhibition balance observed in cortical networks.

#### Network Architecture

The EI network is organized into three distinct layers: an input layer, a recurrent processing layer, and a readout layer. The input layer is responsible for encoding time-dependent sensory signals relevant to the evidence accumulation task. This layer transforms raw sensory data into a format that the network can process efficiently. These encoded inputs are then fed into the recurrent layer, which serves as the computational core of the network.

The recurrent layer is composed of interconnected excitatory and inhibitory units, which capture the dynamics of neural circuits by maintaining a balance between excitation and inhibition. Through these recurrent connections, the network integrates the incoming information over time, allowing it to accumulate evidence continuously from the input stream.

The final readout layer interprets the processed information and produces an output in the form of a decision variable. This abstract decision variable reflects the accumulated evidence, mapping the network’s internal states to a categorical decision or action. The network, through training, learns to optimize the decision-making process by refining the dynamics within the recurrent layer, ultimately producing robust, time-varying outputs aligned with the task’s demands.

#### Input Layer

The evidence accumulation task is a widely used paradigm in both neuroscience and cognitive psychology to investigate the neural and behavioral mechanisms underpinning decision-making processes. In our study, we adapted a task structure akin to the one described by Morcos et al. [29], utilizing a virtual-reality T-maze environment to evaluate decision-making in head-restrained mice. As the mouse progresses through the maze, it is presented with visual cues on both the left and right sides, as depicted in Figure 6D. Upon reaching the critical T-junction, the mouse is required to make a directional choice based on the cumulative number of visual cues from each side, disregarding both the order in which the cues were presented and the position of the final cue. The crux of the task lies in the mouse’s ability to independently count and compare the cues from either side and retain this accumulated evidence until the decision point is reached, preceding any form of reward.

To replicate the complexity of this behavioral task in our conductance-based EI network, we designed the input layer to comprise a population of 100 neurons, segmented into four functionally distinct groups. The first group, containing 25 neurons, encodes the visual cues presented on the left side of the maze, while the second group, also with 25 neurons, encodes the right-side cues. These groups are responsible for accumulating evidence over time, simulating the mouse’s cue-counting process. A third group of 25 neurons becomes active during the recall period, which occurs after the evidence accumulation phase, and is responsible for processing a recall stimulus that prompts decision-making. Finally, the fourth group of 25 neurons generates a steady stream of background noise at a baseline firing rate of 10 Hz, modeling the spontaneous activity typically present in biological networks.

The input stimuli, including both the cues and recall signals, are modeled using Poisson-distributed spike trains at a firing rate of 40 Hz. Each visual cue is represented by a burst of activity lasting 150 milliseconds, followed by a 50-millisecond interval of silence, replicating the temporal dynamics of the task. After the final cue is delivered, a delay period of 1000 milliseconds ensues, during which only the background noise persists. Subsequently, the recall stimulus is presented, cueing the network to generate an output corresponding to the accumulated evidence. This stimulus-response pattern mirrors the behavioral dynamics observed in the mouse as it navigates the virtual maze. These intricate neural representations and task dynamics are further illustrated in Figure 6D.

#### Recurrent Layer

In our spiking neural network model, the recurrent layer is composed of spiking neurons based on a modified version of the Generalized Integrate-and-Fire (GIF) neuron model [22]. This model incorporates one key internal currents: the slow adaptation current. The membrane potential and internal current dynamics are governed by the following equations:

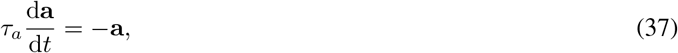

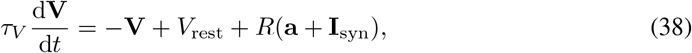

where *τ*_*a*_ represents the time constant of the adaptation current, *τ*_*V*_ = 100 ms is the membrane potential time constant, *V*_rest_ is the resting membrane potential, and *R* = 1.0 Ω is the membrane resistance. The term **I**_syn_ represents the synaptic input from other neurons in the network, and **a** is the spike-triggered adaptation current.

When the membrane potential **V**_*i*_ of the *i*-th neuron reaches the threshold *V*_th_, the neuron generates a spike according to the following update rules:

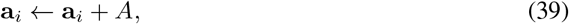

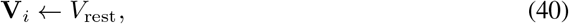

where *A* represents the amplitude of the spike-triggered adaptation current, which is added to the existing adaptation current **a**_*i*_, and the membrane potential **V**_*i*_ is reset to the resting potential *V*_rest_ = 0 mV after each spike.

To simulate the balance of excitation and inhibition observed in biological neural circuits, we populated the recurrent layer with 160 excitatory neurons and 40 inhibitory neurons. This ratio mirrors the natural predominance of excitatory neurons in cortical circuits.

The spike-triggered adaptation current was set with an amplitude *A* = *™* 1.0 mA, creating a negative feedback that regulates neuron firing rates over time. The adaptation time constant *τ*_*a*_ for each neuron was sampled from a uniform distribution 𝒰 [100 ms, 3000 ms] to introduce heterogeneity in neuronal response properties. This variability allows the network to exhibit a rich range of dynamics, facilitating robust temporal integration during the evidence accumulation task.

The external synaptic input **I**_syn_ to each neuron in the network is modeled using a conductance-based exponential synapse model, which captures the dynamics of synaptic transmission in a biologically realistic manner. The total synaptic current **I**_syn,*i*_ received by the *i*-th neuron is described by the following equation:

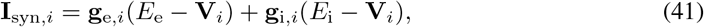

where **g**_e,*i*_ and **g**_i,*i*_ represent the excitatory and inhibitory synaptic conductances for the *i*-th neuron, respectively. The terms *E*_e_ = 3 mV and *E*_i_ = *™* 3 mV are the reversal potentials for excitatory and inhibitory synapses, while **V**_*i*_ is the membrane potential of the neuron. This formulation allows the synaptic current to be modulated by both the conductance changes induced by synaptic input and the difference between the membrane potential and the synaptic reversal potentials, mimicking the biophysical properties of real synapses.

The dynamics of the excitatory and inhibitory conductances are governed by differential equations that describe how these conductances evolve over time:

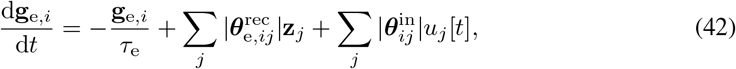

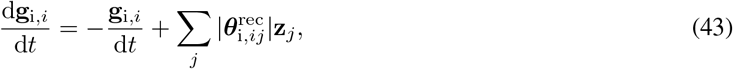

where *τ*_e_ = 10 ms and *τ*_i_ = 10 ms are the time constants for excitatory and inhibitory synapses, respectively, determining how quickly the conductances decay back to zero after synaptic activation. The variables **g**_e,*i*_ and **g**_i,*i*_ represent the excitatory and inhibitory conductances at the *i*-th neuron, which evolve over time based on both recurrent and external synaptic input.

In the above equations, **z**_*j*_ denotes the spike train from the *j*-th presynaptic neuron (either excitatory or inhibitory), with 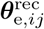 and 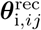 representing the synaptic weights for excitatory and inhibitory connections, respectively, within the recurrent network. Additionally, 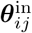 represents the synaptic weights associated with external input (*u*_*j*_[*t*]) to the network. This external input *u*_*j*_[*t*] can represent stimuli from the input layer or other sources outside the network.

The use of absolute values |* | in the synaptic weight terms ensures non-negative conductances, which is a necessary condition for modeling Dale’s principles.

The decay terms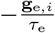 and 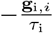 ensure that conductances return to their baseline levels in the absence of ongoing synaptic input, reflecting the transient nature of real synaptic currents.

To inspect the computational graph of the modified GIF network, we also give the discrete description of the model:

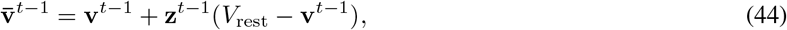

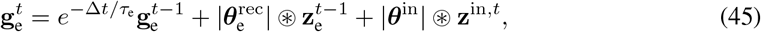

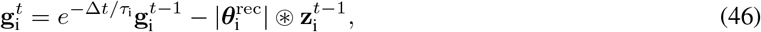

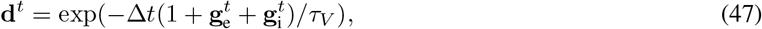

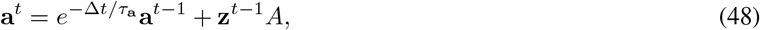

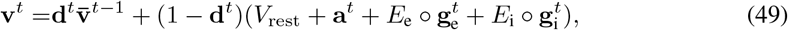

where **d**^*t*^ is an intermediate variable for membrane potential decay, and ∘ is the element-wise multiplication.

#### Readout Layer

All network architectures incorporate a specialized leaky readout mechanism, consisting of dedicated output neurons corresponding to each task-specific label or behavior. These output neurons receive linear projections from the recurrent layer. The temporal evolution of the readout neurons is governed by the following differential equation:

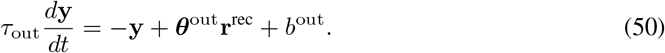

In this equation, **y** represents the activity of output neurons, *τ*_out_ = 10 ms denotes the output neurons’ time constant, ***θ***^out^ signifies the synaptic weight matrix connecting recurrent and output neurons, *b*^out^ is the bias term, and **r**^rec^ denotes spike activities of the recurrent neurons. For computational implementations, a discrete-time approximation of this dynamics is employed:

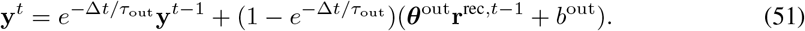

#### Weight Initialization

The readout weights defined above were sampled from a scaled Gaussian distribution, defined as: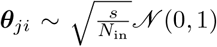, 1), where ***θ***_*ji*_ represents the weight from neuron *i* to neuron *j, N*_in_ denotes the number of afferent (input-providing) neurons, 𝒩 (0, 1) is the standard normal distribution with mean 0 and variance 1, *s* is a scaling factor that allows for control over the initial magnitude of the weights. This initialization schema helps maintain consistent variance of activations and gradients across layers, potentially aiding in training stability and convergence.

For the network with E/I separation, as described in Eq. 42-43, we modified the initialization procedure slightly. While still using the scaled Gaussian distribution, we applied an absolute value operation to ensure all weights were non-negative: 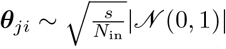.This modification ensures that excitatory remain excitatory and inhibitory connections remain inhibitory, adhering to Dale’s principle often observed in biological networks. The absolute value operation maintains the scale and distribution shape while restricting weights to positive values.

#### Surrogate Gradient Function

Translating membrane potential fluctuations into discrete spike events relies on a Heaviside step function with its transition point at the neuron’s firing threshold *v*_th_. However, this function presents challenges for gradient-based learning algorithms due to its discontinuity at the threshold and lack of meaningful gradients elsewhere. To overcome these limitations and enable effective training of spiking networks, we employed a surrogate gradient approach used in previous works [42].

During the forward pass, we use the Heaviside function to generate the spike:

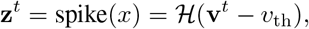

where *x* is used to represent **v**^*t*^ − *v*_th_, and the spiking threshold *v*_th_ = 1 mV.

During the backward pass, we approximated the derivative of the Heaviside step function with the ReLU function:

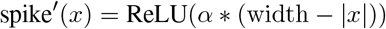

where width = 1.0, and *α* = 0.3. *α* is the parameter that controls the altitude of the gradient, and width is the parameter that controls the width of the gradient. The derivative of the spike function, spike^*′*^(*x*), is a triangular function centered at *x* = 0. It reaches its maximum value of *α* * width at *x* = 0 and decreases linearly to zero at *x* = *±*width. When |*x*| *>* width, spike^*′*^(*x*) = 0.

#### Training Method

The network was trained using the Backpropagation Through Time (BPTT) algorithm, which is well-suited for optimizing spiking neural networks by unrolling their dynamics over time and computing gradients across time steps. To numerically integrate the model’s differential equations, we employed the exponential Euler method, a technique that provides a balance between computational efficiency and accuracy when simulating the continuous-time dynamics of spiking neurons. The integration time step Δ*t* was set to 1 ms.

For gradient-based optimization, we utilized the Adam optimizer [23]. The objective of the training process was to minimize the cross-entropy loss between the network’s predicted output and the target output during the recall phase.

### 5.5 Environment Setting

All evaluations and benchmarks in this study were conducted in a Python 3.10 environment on a system running Windows 11 Home edition. The experiments were executed on a CPU-powered setup, specifically an AMD Ryzen 7 7840HS with Radeon 780M Graphics, clocked at 3.80 GHz. The following software versions were used or compared during the study: `Brian2` (v2.5.1) [33], `BrainPy` (v2.6.0) [39, 40], `Dendritex` (v0.0.1), `braintools` (v0.0.4), `brainstate` (v0.0.2), `brainunit` (v0.0.2), and `JAX` (v0.4.31) [10].

## Data availability

The in *vitro* intra-cellular recording of a cortical pyramidal cell can be obtained in `brain2modelfitting` package. `BrainPy` simulator is available publicly on GitHub at https://github.com/brainpy/BrainPy. The dataset used in this study are open source and publicly available. The evidence accumulation tasks are generated in this study, and can be found in the public available GitHub repository https://github.com/chaoming0625/brainunit-experiments.

## Code availability

`BrainUnit` framework is distributed via the pypi package index (https://pypi.org/project/brainunit/) and is publicly released on GitHub (https://github.com/chaoming0625/brainunit) under the license of Apache License v2.0. Its documentation is hosted on the free documentation hosting platform Read the Docs: https://brainunit.readthedocs.io/. `BrainUnit` can be used in Windows, macOS, and Linux operating systems. The code to reproduce the experimental evaluations in this study is publicly available from the following GitHub repository: https://github.com/chaoming0625/brainunit-experiments. `BrainUnit` has been seamlessly integrated into the ecosystem of brain dynamics programming, which can be obtained online through: https://ecosystem-for-brain-dynamics.readthedocs.io/.

## Acknowledgments

This work was supported by the Science and Technology Innovation 2030-Brain Science and Braininspired Intelligence Project (No. 2021ZD0200204, SW), the Young Scientists Fund of the National Natural Science Foundation of China (No. 3240070449, CMW), the China Postdoctoral Science Foundation (No. 2024M750076, CMW), and the Postdoctoral Fellowship Program of CPSF (No. GZC20230106, CMW).

## Author contributions

Conceptualization and Methodology: CMW. Software and Investigation: CWM, SCH, SWL. Analysis: CMW. Visualization: CMW, SCH. Writing: CMW. Writing (Review & Editing): CWM, SCH, SWL, YXH, SW. Funding acquisition: SW, CMW. Resources: YXH, SW. Supervision: CMW, SW.

## Competing interests

Authors declare that they have no competing interests.

## Supplementary Information

### A Extending Units

`BrainUnit` is a powerful library designed to facilitate the flexible definition, customization, and extension of physical units in scientific computing. This flexibility manifests across multiple aspects, including dimensions, metric scales, and base factors (as illustrated in Figure 1). The library’s architecture allows users to create and manipulate units with precision and ease, catering to a wide range of scientific and engineering applications.

#### A.1 Fundamental Components of a Unit

To instantiate a new `Unit` object, the following parameters need to be specified:

- **dim**: A `Dimension` instance that defines the unit’s physical dimensions. These are created using the `brainunit.get_or_create_dimension` function or instantiating the `brainunit. Dimension` class.
- **base**: A float value specifying the base factor of the unit. By default, this is set to 10, adhering to the convention of SI units. However, it can be customized for different unit systems.
- **scale**: An integer specifying the exponent for the scale factor. The default value is 0, representing the base unit.
- **name**: A string representing the full name of the unit (e.g., `“volt”`).
- **dispname**: A string for the display name or symbol of the unit (e.g., `“V”`).

#### A.2 Creating New Units

`BrainUnit` offers multiple methods for creating new units, each serving distinct purposes:

**Direct Instantiation of the** `Unit` **Class**. The most fundamental method involves directly instantiating the `Unit` class:

**Listing S1:**
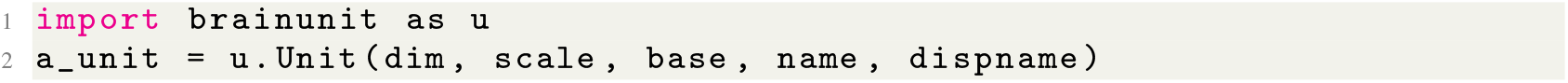
Creating a New Unit by Direct Instantiation

This method is primarily used in internal computations, especially when composing multiple units during complex calculations. For instance, `u.mS / u.cm ** 2` would create a unit with the name `“msiemens / cmeter2”`. Units created this way are not automatically registered as standard units.

**Using the** `Unit.create()` **Method**. For creating standard units that can be used in further compositions, the `Unit.create()` method is preferred:

**Listing S2:**
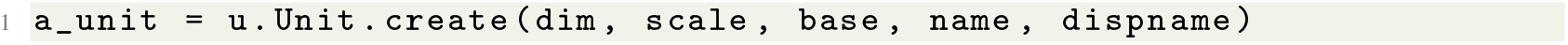
Creating a New Standard Unit

This method registers the newly created unit as a standard unit within the `BrainUnit` system.

**Creating Scaled Units**. To create scaled versions of existing units, `BrainUnit` provides the `Unit. create_scaled_unit()` method:

**Listing S3:**
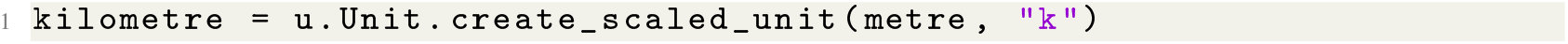
Creating a Scaled Unit

This method takes two parameters:

- **baseunit**: The original unit to be scaled (e.g., `metre`).
- **scalefactor**: A string specifying the scaling prefix (e.g., `“k”` for kilo, `“m”` for milli). Available prefixes are listed in Table S1.

#### A.3 Creating New Dimensions

Creating new units often requires defining new dimensions. `BrainUnit` offers two methods for this purpose:

**Direct Instantiation of the** `Dimension` **Class**. The most fundamental method involves directly instantiating the `Dimension` class:

**Listing S4:**
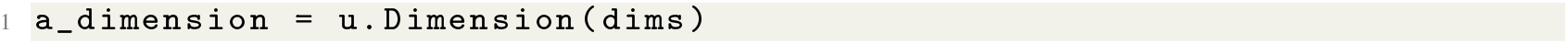
Creating a New Dimension by Direct Instantiation

Here, `dims` is a vector of seven integers representing the exponents of the seven basic SI units (refer to `brainunit.Dimension` for Automatic Unit Conversion for details).

**Using the** `get_or_create_dimension()` **Function**. A more intuitive method involves using the `brainunit.get_or_create_dimension` function:

**Listing S5:**
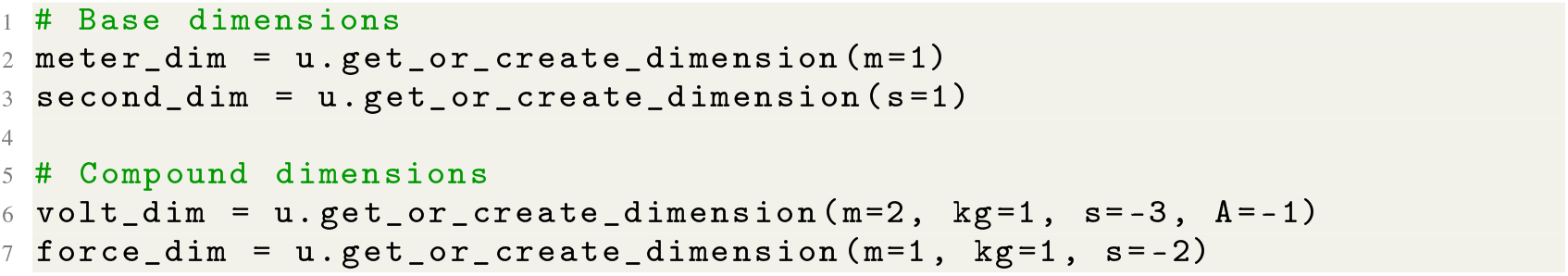
Creating Dimensions with `get_or_create_dimension()` Function.

This function accepts both keyword arguments (e.g., `m`, `kg`) and positional arguments corresponding to the seven fundamental SI dimensions.

#### A.4 Customizing the Base Factor

While `BrainUnit` defaults to base 10 for SI units, it allows for customization to accommodate different unit systems:

**Listing S6:**
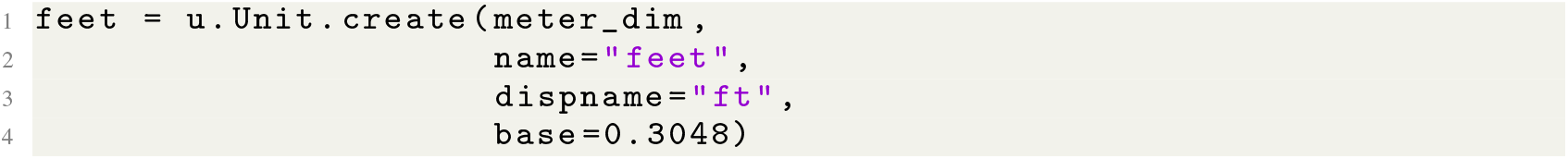
Defining a `feet` Unit with a Different Base Factor

This flexibility enables the creation of units in various systems, such as U.S. customary units or binary systems (base 2).

#### A.5 Specifying Scale Exponents

`BrainUnit` provides a range of predefined scale prefixes for creating scaled units efficiently. These prefixes, listed in Table S1, allow for easy creation of units across different orders of magnitude.

By leveraging these components and methods, `BrainUnit` provides a comprehensive and flexible system for defining, customizing, and working with physical units in scientific computing applications.

### B Supplementary Tables

**Table S1:**
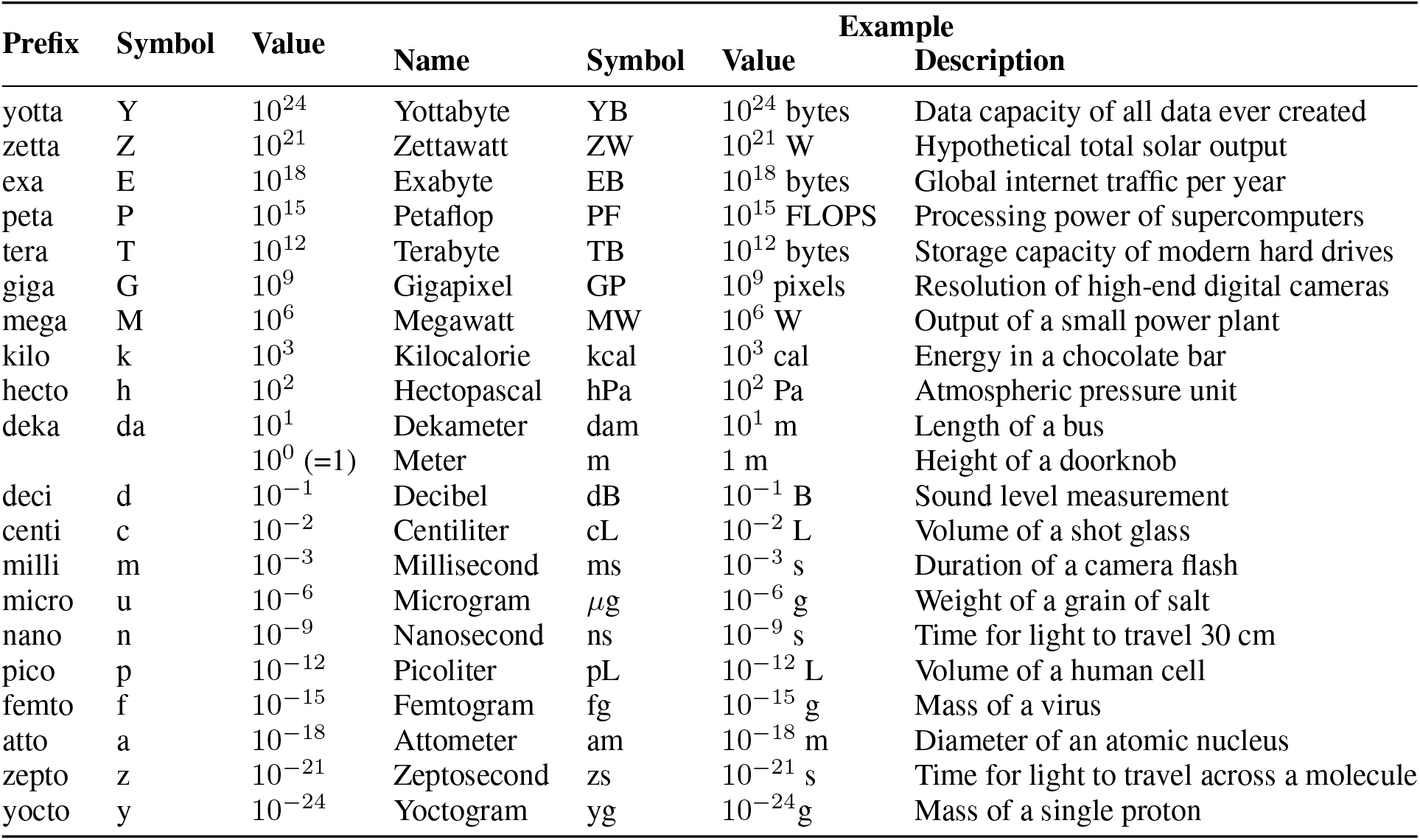
Metric Prefixes, Symbols, and Examples

**Table S2:**
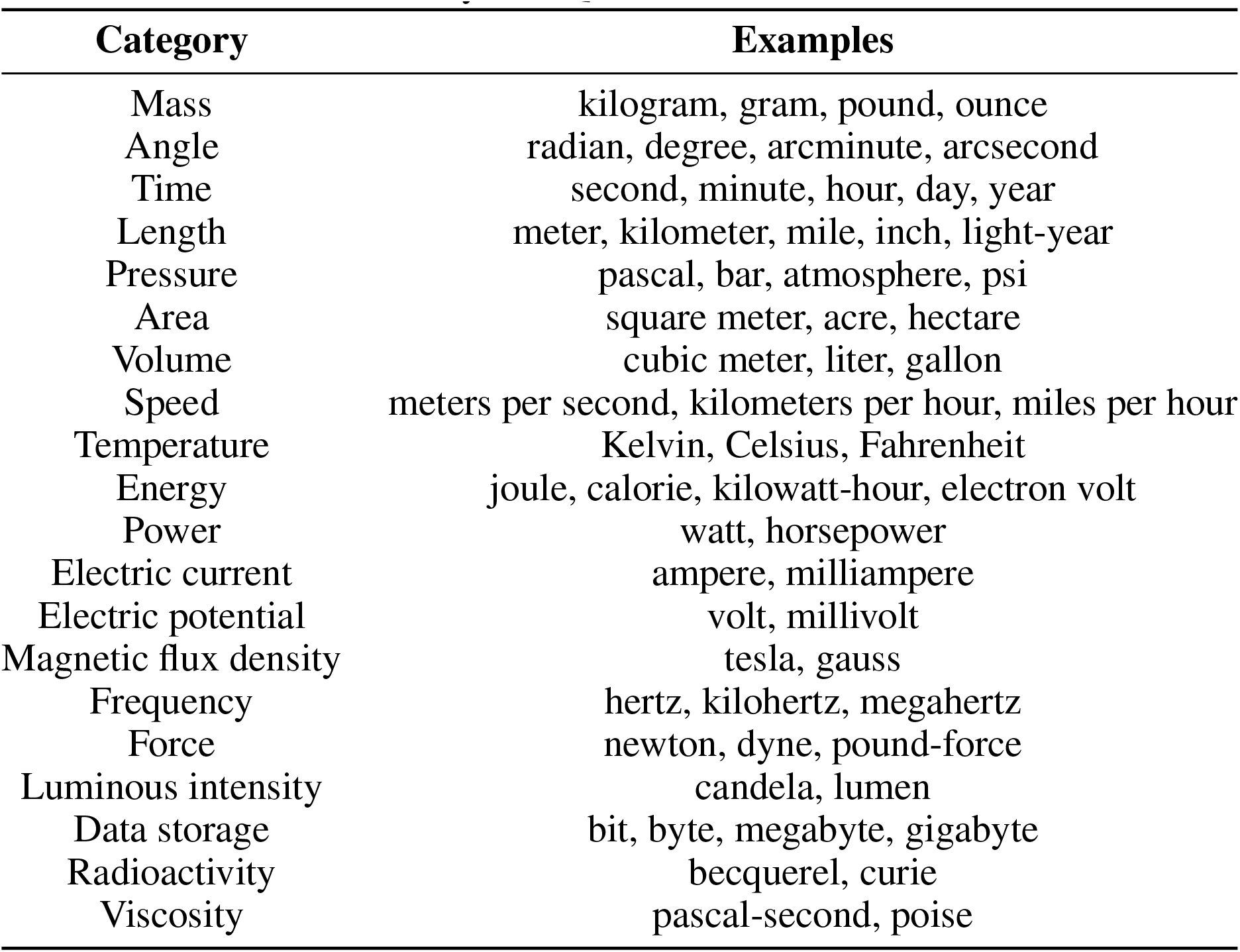
Physical Quantities and Their Units

## Notes

### Competing Interest Statement

The authors have declared no competing interest.

### Summary of Updates

Change the title. Add new texts.

